# Contrasting leaf thickness & saturated water content explain diverse structural/physiological properties of arid species

**DOI:** 10.1101/2022.11.02.514850

**Authors:** Byron B. Lamont, Heather C. Lamont

## Abstract

Although they account for many thousands of the world’s flowering plants, little is known about the physical/chemical properties of leaf succulents. Eight species in the Namib Desert, South Africa were assessed for leaf area (*A*) and thickness (*z*), saturated (*Q*) and dry mass, relative volume of air (*F*_a_), intrinsic water-us efficiency (δ^13^C), and N, P and (Na+K) contents. As water-storage capacity is a function of *Q*_v_ and *z*, this means *Q*/*A* (= *Q*_v_ •*z*) is an ideal index of succulence compared with specific-leaf-area and other indices that highlight mass rather than volume. Specific gravities have a different relationship with *F*_a_ of sclerophyll-mesophylls: rising succulence infers decreasing air content replaced by water rather than dry matter. The trend among succulent species from Argentina/Spain added to our data was characterized by *Q*/*A* exceeding 1 mg water/mm^2^ leaf whose overall slope was ten times that for co-occurring sclerophyll-mesophyll species. (Na+K), N and P concentrations varied on a dry-matter, but not water-volume, basis. *W*_i_ relationships were essentially functions of variations in *z* (transpiration-resistance) and increased metabolic efficiency. We conclude that *z* and *Q*_v_ are keys to the special physiological properties of succulent leaves. Including succulents would force many current monotonic relationships to dichotomize.

**Highlight:** The key to understanding leaf succulents is their high volumetric water content and thick leaves. These explain their superior water-use-efficiency and contrasting relationships with other variables compared with temperate species.

## Introduction

All leaves store water, even if only over night. At the other extreme, succulent leaves retain water for weeks or months even when severed from their supporting root systems (von Willert & Brinckman, 1986; Lamont & Lamont, 2000). Prolonged storage during drought is an alternative mechanism to continuous contact with soil moisture as a source of water for use by the plant (Milton *et al*., 1997) and these correspond to a dichotomy in the penetration of the soil profile by the root systems (Kutschera *et al*., 1997; Lamont & Lamont, 2000). There is much interest in the structural versus functional properties of leaves. At one level, there are attempts to relate leaf structure to their environment (Lamont *et al*., 2002; Wright & Westoby 2002; Grubb et al. 2015). At another, the aim is to identify the basic components of leaf structure and see how they vary between species and each other (Niinemets, 1999; Pyankov *et al*., 1999; Roderick *et al*., 1999b; Wang et al. 2022). Few of these studies have included leaf succulents. This implies bias; e.g., there are 5,000 species just in the leaf-succulent family, Mesembryanthemaceae (Milton *et al*., 1997). Exceptions include Vendramini *et al*. (2002), who studied 13 leaf succulents among 77 Argentine species, and Grubb *et al*. (2015) who included 11 species that they considered leaf succulents/semi-succulents among 38 arid Spanish species. It is usually anticipated that generalities determined for non-succulents will not apply to succulents (Roderick *et al*., 1999c). In concluding that leaf dry-matter content (*D*_m_) was a useful functional trait, Wilson *et al*. (1999) noted that it was not clear how relevant it would be for dry climates with many succulents. Vendramini *et al*. (2002) showed that some relationships between leaf properties depended on whether succulents were included in the sample, while Grubb et al. (2015) noted that the physical properties of their leaf succulents were quite different from the non-succulents.

We studied eight species, varying from semi-sclerophyllous to highly succulent, in the southern Namib Desert, South Africa, with five species having a saturated water content (*Q*_v_) > 90% and thus fitted the leaf succulent class (Vendramini *et al*., 2002). The vegetation is part of the succulent karoo that Grubb et al. (2015) specifically noted would form a useful comparison with their study in semi-arid Spain. Our previous paper explored the concept of ‘utilizable’ water, and showed that the range for these species was a water content (*Q*_m_) at full turgor (saturated) of 72−93%, based on a critically low level of relative fluorescence (Lamont and Lamont, 2000).

Here, we examine, theoretically and empirically, how three standard measures of succulence relate to their possible underlying properties, such as leaf thickness (*z*), and dry matter and saturated water contents on the basis of mass (*D*_m_ and *Q*_m_) and volume (*D*_v_ and *Q*_v_). We were aware of the problem of comparing confounded variables as they will be correlated by definition (Williams et al. 2022), unless they remain constant in a particular study. Thus we note, for example, that the type of relationship of *D*_v_ with specific leaf area [SLA = 1/(*D*_v_•z)], proposed as of potential interest by Wilson *et al*. (1999), is already mathematically prescribed. When undertaken here, the aim was to determine which of the various components best explained the relationship. Logically, mass-related indices are more likely to be related to nutrient-storage properties, and volume-related indices to water-storage properties. Because Roderick *et al*. (1999b) highlighted the need to recognize air as a leaf component, we examined how the relationship of air on a volume basis (*F*_a_) varied with water (*F*_Q_ = *Q*_v_) and dry matter (*F*_D_). This analysis first required us to develop a technique to determine the volume of dry matter that usually remains unknown.

The volume (v) of a leaf is made up of the product of the projected area (*A*) and mean thickness (*z*) (Lamont et al. 2015). Contributors to volume are dry matter, water and air. The leaf’s turgid mass consists of its dry mass (*D*) plus water content (*Q*). Dry mass and water content can be related to turgid mass (*D*_m_ and *Q*_m_ respectively). They can also be related to turgid volume (*D*_v_ and *Q*_v_ respectively). Specific leaf area (SLA) is given by *A*/*D*_m_ and corresponds to the inverse product of *D*_v_ and *z* (Witkowski & Lamont, 1991). Increasing succulence can be expected to involve an increase in *Q*_v_ with a decrease in air (*F*_a_) and volume of dry mass (*F*_D_), and thicker leaves (*z*) imply greater succulence. Three standard indices of leaf succulence have been used historically:

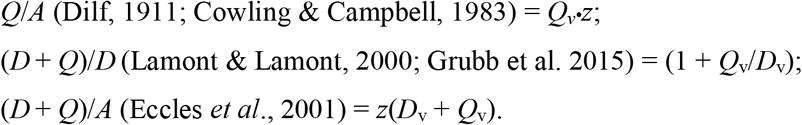

Our aim was to determine for our eight species which of their components most affected these indices of leaf succulence, and which index best reflected the water-storing capacity of a leaf.

While not necessarily high in inorganic cations, especially Na^+^ and K^+^ (a property of halophytic succulents), these might have an osmotic function in promoting expansion of the cell vacuole, and were examined here. Roderick *et al*. (1999b) also pointed to the functional value of nitrogen content (N) per unit water, so we explore how N on a volume basis (N_v_) varied with the indices of succulence, *Q*_v_ and *z*. N/*A* and P/*A* (nitrogen and phosphorus on an area basis) should indicate the assimilatory and metabolic potential of a leaf and have been shown to increase with increasing aridity (Lamont *et al*. 2002) – how then do these vary with structural properties? On the other hand, since increasing thickness in succulents, as a response to aridity, is due to the accumulation of water-storing mesophyll rather than photosynthetic surface tissues (Nobel *et al*., 1994), we might expect N/*A* and P/*A* to be unaffected. Finally, there has been much research on how the carbon isotope discrimination ratio (δ^13^C) varies with the type of photosynthesis and environment but little with leaf properties (e.g., Hanba *et al*. 1999). We therefore examine if there is any relationship between δ^13^C (as intrinsic water-use efficiency, *W*_i_) and SLA, *D*_v/m_, *z, Q*_A_, N/*A* and P/*A*. We would expect *W*_i_ to increase as succulence increased (Teeri *et al*., 1981; Maxwell *et al*. 1997).

## Materials and Methods

Leaves were collected from eight species growing at Groenriviersond, 500 km north of Cape Town, South Africa (30º 51’ S, 17º 34’ E). The site lies in the southern portion of the Namib Desert. Rainfall was 79 mm in the year of the study (1998) although fog and dew are regular occurrences. The vegetation is part of the succulent karoo and consists of clumps of creepers to woody shrubs up to 2 m tall (Eccles *et al*. 2001). The soil is red aeolian sand overlying an impenetrable silcrete hardpan at about 2 m depth. The species were selected to cover the full range of apparent water-holding properties at the site and consisted of *Pteronia onobromoides* (Asteraceae), *Salvia lanceolata* (Lamiaceae), *Eriocephalus africanus* (Asteraceae), *Stoeberia utilis* (Mesembryanthemaceae), *Ruschia fugitans* (Mesembryanthemaceae), *Zygophyllum morgsana* (Zygophyllaceae), *Othonna cylindrica* (Asteraceae) and *Senecio* aff. *sarcoides* (Asteraceae). Leaves of all species were iso(bi)lateral and sessile. Accepting 90% saturated water content on a mass basis (*Q*_m_) as the critical level for designation as a succulent (Gibson, 1996 in Vendramini *et al*., 2002), then the first three species were non-succulent (75–84%) and the last five were succulent (92–97%). On a preferred volume basis, *Q*_v_, two species were in the range 40−50%, three were 60−70% and three were 80−95%. No equivalent critical level is recognized at present for the three indices of succulence noted in the Introduction. More importantly, the properties of the eight species studied here formed a well-defined gradient comparable with, and oftern exeeding, values previously obtained for the same indices (Table 1). Nomenclature is given in Eccles *et al*. (1999) and, from hereon, only the genus names are used.

**Table 1.**
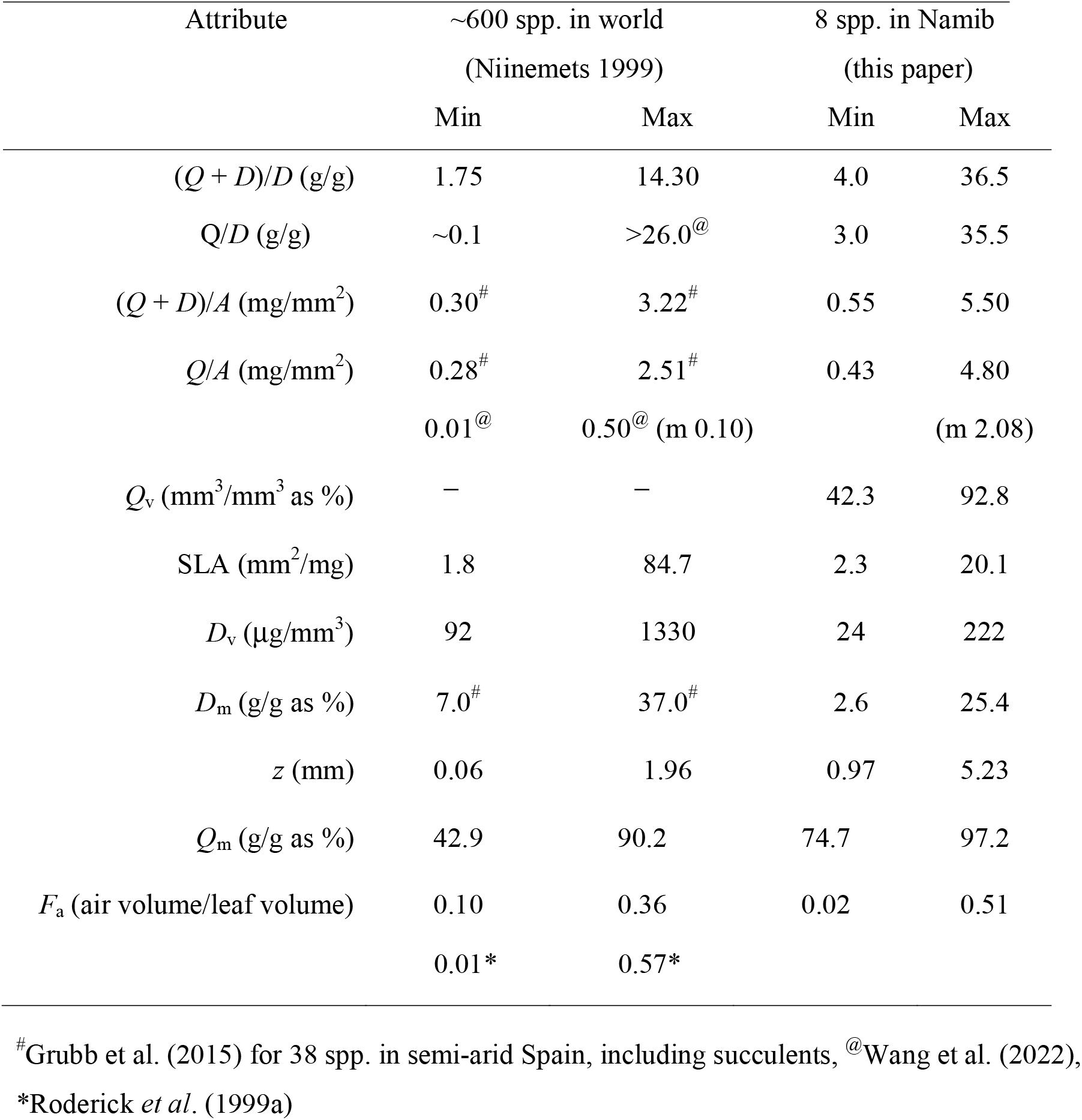
Limit of physical properties of leaves worldwide compared with the eight species studied here. The first three attributes are common indices of leaf water-storage capacity. m = mean.

| Attribute | ~600 spp. in world<br>(Niinemets 1999) |  | 8 spp. in Namib<br>(this paper) |  |
| --- | --- | --- | --- | --- |
|  | Min | Max | Min | Max |
| $(Q + D)/D$ (g/g) | 1.75 | 14.30 | 4.0 | 36.5 |
| $Q/D$ (g/g) | ~0.1 | >26.0<sup>@</sup> | 3.0 | 35.5 |
| $(Q + D)/A$ (mg/mm<sup>2</sup>) | 0.30<sup>#</sup> | 3.22<sup>#</sup> | 0.55 | 5.50 |
| $Q/A$ (mg/mm<sup>2</sup>) | 0.28<sup>#</sup> | 2.51<sup>#</sup> | 0.43 | 4.80 |
|  | 0.01<sup>@</sup> | 0.50<sup>@</sup> (m 0.10) |  | (m 2.08) |
| $Q_v$ (mm<sup>3</sup>/mm<sup>3</sup> as %) | — | — | 42.3 | 92.8 |
| SLA (mm<sup>2</sup>/mg) | 1.8 | 84.7 | 2.3 | 20.1 |
| $D_v$ (μg/mm<sup>3</sup>) | 92 | 1330 | 24 | 222 |
| $D_m$ (g/g as %) | 7.0<sup>#</sup> | 37.0<sup>#</sup> | 2.6 | 25.4 |
| $z$ (mm) | 0.06 | 1.96 | 0.97 | 5.23 |
| $Q_m$ (g/g as %) | 42.9 | 90.2 | 74.7 | 97.2 |
| $F_a$ (air volume/leaf volume) | 0.10 | 0.36 | 0.02 | 0.51 |
|  | 0.01* | 0.57* |  |  |

<sup>#</sup>Grubb et al. (2015) for 38 spp. in semi-arid Spain, including succulents, <sup>@</sup>Wang et al. (2022),
\*Roderick *et al.* (1999a)

Current season’s mature twigs (100–150 mm long) were removed from side branches of 6–8 plants of each species by cutting under water predawn. They were kept with their ends in water at 17.5–20.5ºC and covered with plastic bags for 1–4 days in the laboratory to ensure full hydration. They were then recut under water and their pressure-volume relations determined following the protocol of Radford & Lamont (1992). The balancing pressure was achieved with a digital pressure chamber, model 1003, PMS Instruments, Corvallis, OR, USA. Wet weight values of twigs were extrapolated to *Ψ* = 0 to obtain turgid (saturated) mass. Ten mature, full-sized leaves were removed from other stems, and these plus the original twigs used were weighed, frozen at −16ºC to rupture the cells and hasten drying, dried at 72ºC for 48 h and reweighed. From this, turgid mass of the twigs was used to obtain leaf turgid mass (60–95% of total mass for individual twigs). Midpoint thickness of 10 leaves from 3 plants was determined with callipers. Projected area (*A*) was obtained by placing 30 leaves or more diagonally on the conveyor belt of an area meter (Li-Cor 3000, Lincoln, NK, USA). Adjustments were made for the shape of leaves and their volume (*v*) determined geometrically (Lamont et al. 2015): five were cylindrical (*v* = π/4*z*•*A*) where *z* was diameter, two were laminate (*v* = *z*•*A* where *z* was thickness) and one was subulate (*v* = mean *z*•*A*), all lacking midribs. SLA [*A*/*D* = 1/(*D*_m_•*z*), where D = dry leaf mass and *D*_m_ = dry leaf density on a mass basis, Witkowski and Lamont 1991] was adjusted for leaf shape in the same way, and *D*_v_ and *Q*_v_ (dry matter and water mass per unit leaf volume) were based on these measurements.

Volume of dry matter was determined by removing all, and only, mature leaves from six twigs, bulking and macerating after oven-drying as above to pass through a 1.1 mm mesh, then twice through a 0.3 mm mesh. The powder was then moistened with a wetting agent (1% Tween 20) in distilled water to form a thick paste. A cork borer (internal diameter 3.58 mm) was pushed against the paste, to produce an initial firm cylinder 20–40 mm in length. It was then placed on a paper tissue over plastic sheeting on a fibrocement base. A iron rod of diameter 3.50 mm was pushed into the borer and tapped with a small hammer about 30 times, until water no longer squeezed out of the bottom. The pressure applied was up to 5.1 kg/cm^2^ but it was usually about 2.1 kg/cm^2^. The cylinder of compressed paste was forced out with the rod, and the ends cut with a razor blade as required to produce a perfect cylinder and its length determined with callipers. 3–5 cylinders were obtained per species. They were dried at 65ºC for 40 h and kept in a desiccator until weighing. Knowing *D*_v_ (*D*/*V*) and volume of dry matter per unit dry mass (*V*_D_/*D*), the contribution of dry matter volume (essentially cell walls, but including protein and solutes) to total volume (*V*_D_/*V* = *F*_D_) and air, *F*_a_ = (1 – *F*_D_) were calculated. Thus, colloidal protein and soluble substances were put with structure rather than cytoplasm or vacuole when estimating volume fractions (Roderick *et al*. 1999b). Some solutes may not have been adsorbed by the cell-wall components, but, even if not, their mass accounted for < 0.01% of dry matter according to our estimates. Methods for specific gravity (*ρ*) followed Roderick *et al*. (1999a).

Since Wang et al. (2022) demonstrated a significant relationship between SLA and *Q*/*D* for thousands of species that did not appear to include leaf succulents, we decided to compare *Q*/*D* with SLA (*A*/*D*), whose slope produces *Q*/*A* – our preferred index of succulence, with data sets that include succulents: ours, Vendramini et al. (2002) and Grubb et al. (2015). This provided 34 species that the researchers considered to be succulents, and 69 non-succulents. We converted the *Q*/(*Q* + *D*) values of Vendramini et al. (2002) to *Q*/*D* by a) inverting the value, b) subtracting 1, and c) re-inverting the new value. Inspection of our data indicated that *Q*/*A* for the succulent species was ≥ 1mg of water per 1 mm^2^ of leaf surface and all 21 species in the data set conforming with this range were previously considered to be succulents; so we set this as the critical value defining succulents. Two of our species were in the range 0.5–1.0 mg/mm^2^ and 10 of 27 species in this range were previously considered succulents; so we assigned these as semi-succulents. Two of 34 species in the data set were < 0.5 mg/mm^2^ and their SLA was < 10 mm^2^/mg (LMA = 100 µg/mm^2^) that seemed reasonable boundaries for sclerophylly (Wright et al. 2003) and also points to how LMA fails as a good index of sclerophylly when succulents are included in the study. None of 33 species in the *Q*/*A* < 0.5 mg/mm^2^ class with indeterminate SLA were previously considered succulent and we suggest that these fit the mesophyll class of leaf structure, of which sclerophylly forms a subset. Since the succulent and sclerophyll-mesophyll classes clearly formed different patterns, we determined the best fit curves for both and added them to the graph.

Total P, K and Na were obtained on 0.5 g samples of the cylinders by digesting with nitric acid and analysing by inductively coupled plasma spectroscopy (WAITE Labs, Adelaide, South Australia). Total N was determined similarly but using standard Kjeldahl digestion and titration. Protein content was taken as 6.25N and N determined on a volume basis (N_v_) based on its mass contribution to *D*_v_. Dry leaves from the physical analyses were subsampled and 0.5 mg analysed in duplicate for the stable isotopes ^12^C and ^13^C and the standardized ratio, δ^13^C, determined (Farquhar *et al*., 1982). This was converted to intrinsic water-use efficiency (*W*_i_) using the equations given in Swanborough *et al*. (2003). A mass spectrometer (Finnigan MATT 252) was used after tissue combustion in a Fisons CHN analyser under continuous flow and compared against the PDB standard (Ehleringer & Osmond, 1989).

Pairwise comparisons of attributes of interest were made using standard best-fit curves (linear, exponential, logarithmic, power) provided by the Cricket Graph III graphing program (Computer Associates, USA). Where no directional relationship was expected, a 2-tailed test was used with *P* = 0.05 for *r* = 0.706, *P* = 0.01 for *r* = 0.834 and *P* = 0.001 for *r* = 0.925. Where a relationship was expected from theory, a 1-tailed test was used with *P* = 0.05 for *r* = 0.621, *P* = 0.01 for *r* = 0.700 and *P* = 0.001 for *r* = 0.788. Solid or dotted lines were drawn to indicate significant relationships and dashed lines when the trends were only significant at P < 0.06.

## Results and Discussion

### Indices of succulence

Compared with other species worldwide, some of our species showed exceptionally high levels of the three indices of succulence, leaf thickness (*z*) and saturated water content on a mass basis (*Q*_m_), and low levels of SLA and dry density on a saturated mass basis (*D*_m_), and an air volume content (*F*_a_) that exceeded the range recorded by Niinemets (1999) (Table 1). Including our Q/A values in theThere were strong positive linear correlations between saturated water content on a volume basis (*Q*_v_) and *z* with two of the indices of succulence, (*Q* + *D*)/*A* and *Q*/*A*, but no (curvi)linear fit with the mass-based index, (*Q* + *D*)/*D* (Fig. 1a,b). (*Q* + *D*)/*A* is equivalent to (*Qv•z* + *D*_v_*•z*) and *Q*/*A* to *Qv•z* so that their significant relationship with *Q*_v_ and *z* is no surprise as they are correlated by definition (Williams et al 2022). Garnier & Laurent (1994) reported a positive slope of 0.5 between (*D* + *Q*)/*A* (= 1 + *Q*_v_/*D*_v_) and z for grasses while ours was 1.2, no doubt because of the additive effect of the associated increasing *Q*_v_ with rising succulence. Their *z* values stopped 750 μm before ours started and ours continued until 5230 μm. Roderick *et al*. (1999a) also obtained a positive relationship between *Q*/*A* and *z* for non-succulents (their *z* values stopped before ours started) but their slope was lower (0.4 vs 1.0) for the same reasons.

**Fig. 1.**
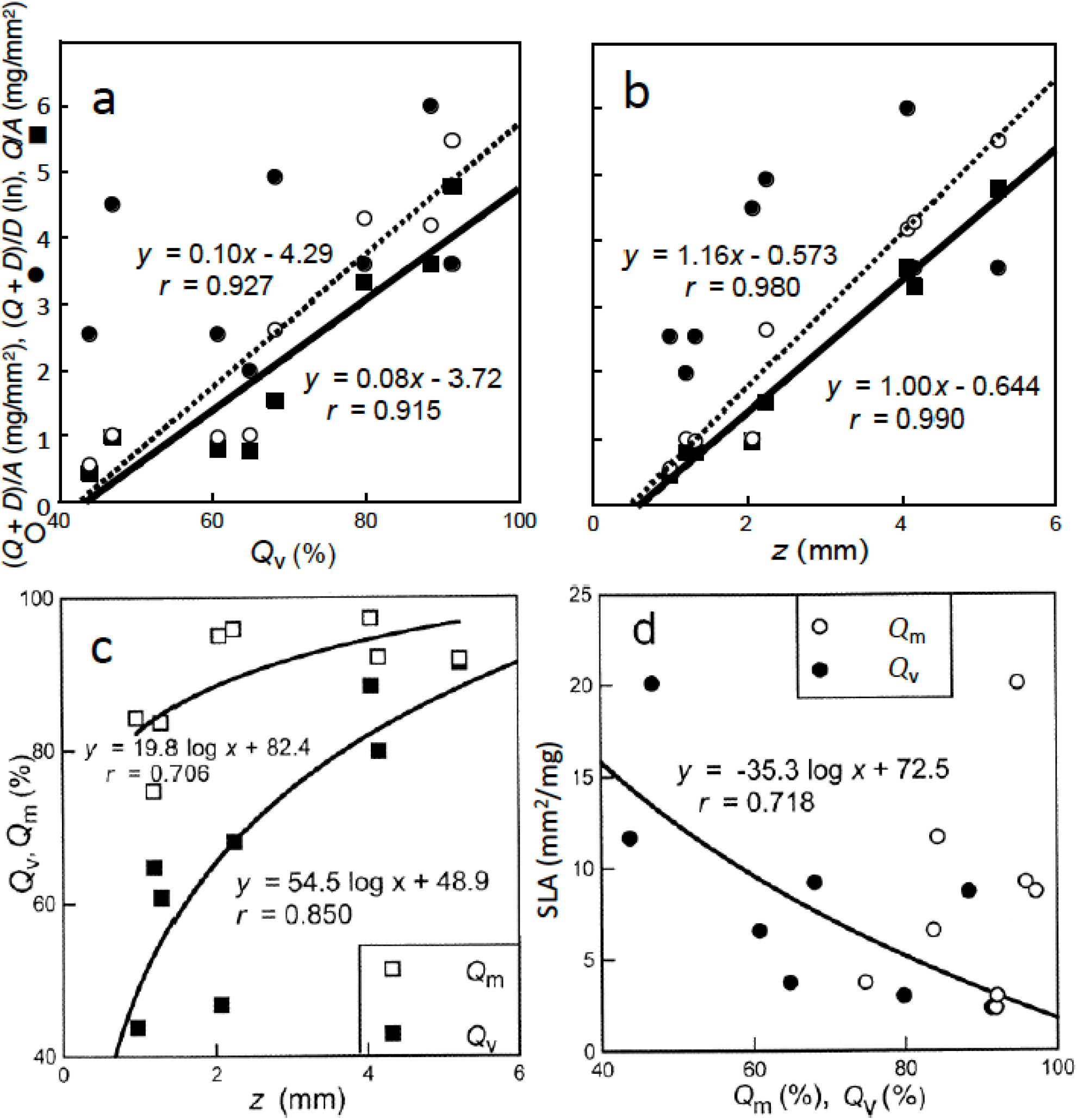
Three indices of succulence, (*Q* + *D*)/*D*, (*Q* + *D*)/*A* and *Q*/*A*, plotted against a) *Q*_v_, and b) *z*; c) *Q*_m_ and *Q*_v_ plotted against *z*; and d) SLA plotted against *Q*_m_ and *Q*_v_. Best-fit curves for (*Q* + *D*)/*A* given by dotted lines and *Q*/*A* by solid lines.

Choice of index of succulence made a difference of up to four places in the rankings among our eight species. (*Q* + *D*)/*D*) cannot be adequate as it ignores the third structural component of cells, air space (*F*_a_) (Roderick *et al*., 1999b). We showed that a *Q*_m_ of 75–97% can be accompanied by an air space of 2–51%, such that there is no relationship between *Q*_m_ and *Q*_v_ (Fig. 1d). Since succulence is clearly a concept based on high water storage per unit leaf volume, rather than mass, and thicker leaves with equal saturated water content are considered more succulent from a textural perspective, this makes *Q*/*A* the obvious choice as the *best index of succulence*, for it is the product of water storage capacity on a unit volume basis, *Q*_v_, and leaf thickness, *z*. Interestingly, there was a clear logarithmic relationship between *Q*_v_ and *z* with *Q*_v_ gradually approaching 100% as leaves continued to thicken. A negative relationship was obtained by Roderick *et al*. (1999a,c) who proposed it as general rule but it cannot apply to leaf succulents where increased water storage capacity is clearly associated with increased z. Our values for *A*/*Q* would force the slope of 0.97, obtained by Wang et al. (2022) for several thousand species worldwide, to fall convexly to a mean of 0.05 (Table 1) as Q increased.

### Water vs air

As *Q*_v_ (= *F*_a_) increased from 42% to 93%, the fraction of air space, *F*_a_, decreased from 51% to 2% in a strongly linear manner, while the fraction of dry matter volume, *F*_D_, varied little at ∼6.4% (Fig. 2a). Thus, saturated water content on a volume basis increases at the expense of air not dry matter. This means for an increase in *Q*_v_ from 40% to 90%, cell wall area increases by 33% so that their thickness is reduced by 25% (Fig. 2c). Thus, the mesophyll of succulents has relatively large, thin-walled cells, a protoplast engorged with water, and tiny intercellular air spaces that would limit diffusion in the apoplast. Roderick *et al*. (1999b) is correct in emphasising the need to recognize *F*_a_ as the third fundamental property of leaves. Since its contribution to leaf volume varied by 50% in our species, *Q*_m_ and *Q*_v_ were completely uncorrelated. This means that using *Q*_m_ as a surrogate for *Q*_v_ or *D*_v_, as undertaken in some functional ecology studies (Weiher *et al*., 1999, Wang et al. (2022), could be quite misleading. Thus, it is difficult to advocate substituting *Q*_m_ (or *D*_m_) for *Q*_v_ (or *D*_v_) on theoretical grounds as they lack a structural basis (the presence of air is ignored). On practical grounds, the only additional measurement needed to estimate volume is *z*, which appears to be an even more fundamental functional property than both (see below).

**Fig. 2.**
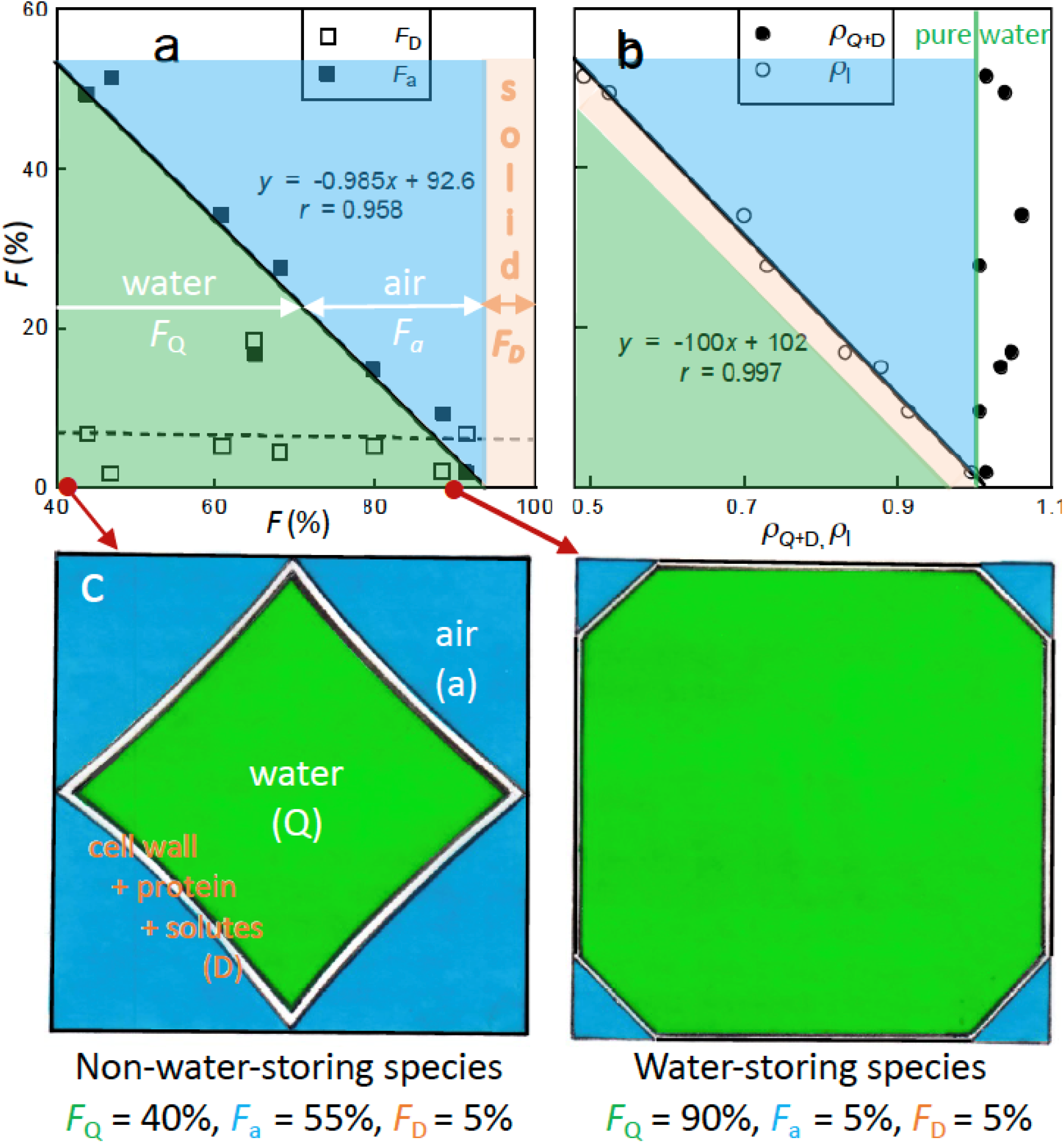
Volume fractions of major cell components (*F*). a) *F*_D_ (orange area) and *F*_a_ (blue) plotted against *F*_Q_ (= *Q*_v_, green), *F*_D_ and *F*_a_; b) *F*_Q_ and *F*_D_ plotted against *ρ*_Q+D_ and *ρ*_l_ (note *ρ*_Q+D_ is slightly denser than pure water); c) Model of relative volume of cell components from data in Figs. 1, 2, 4; left cell based on *F*_Q_ = 40% (≡ *Salvia*), right on *F*_Q_ = 90% (≡ *Ruschia*). Green *Q*_v_, blue *F*_a_, white = cell wall (*F*_D_ – *F*_protein_), inner line = *F*_protein_. Because of the colloidal properties of protoplasm its volume cannot be defined, so it is separated out on the inner wall as *F*_protein_, and all water is treated as in the vacuole. Model assumes an average cell of both occupies the same space; in practice, the more succulent species has larger cells.

Specific gravity for *Q* + *D* (*ρ*_Q+D_) (i.e., omitting air spaces) varied from 1.007 to 1.034 with a mean of 1.025 ± 0.020 (SD) and showed no relationship with *F*_a_ (Fig. 2b). Specific gravity for the whole leaf, *ρ*_l_, (i.e., including air spaces) ranged 0.492–0.995 with a mean of 0.763 ± 0.184 and had an almost perfect linear relationship with *F*_a_. Our *ρ*_Q+D_ and *ρ*_l_ values are in the same range as those for non-succulents (Roderick *et al*., 1999a,b). But *ρ*_Q+D_ showed no relationship with the excluded fractional air space (*F*_a_) (positive for Roderick *et al*., 1999a) and *ρ*_l_ showed a strong negative relationship with *F*_a_ (positive for Roderick *et al*., 1999a). This points to a basic difference between the structure of mesophytes-sclerophylls and succulents: *ρ* rises through an increase in dry matter density (*D*_v_) accompanied by a decrease in water content (*Q*_v_) among non-succulents, whereas *ρ* rises through an increase in water at the expense of air in succulents. Volume of dry matter (*F*_D_) plays no role in interpreting the wide variation in *F*_a_ and *Q*_v_ among our species as it varies little (Fig. 2b).

### Relationships with SLA

There was no relationship between the three indices of succulence and SLA (Fig. S1). However, SLA declined logarthmically with increase in *Q*_v_, a component of *Q*/*A* (Fig. 1d). *Q*_v_ (and to a lesser extent *Q*_m_) increased at a decreasing rate as *z* increased (Fig. 1c). The lack of a (negative) correlation between *Q*/*A* (= *Qv•z*) and SLA [= 1/(*D*_v_•z)], unexpected because of their common but inversely related *z*, must therefore be due to the erratic relationship between *Q*_v_ and *D*_v_ (Fig. S2). Note that *z* also increases as sclerophylly (the opposite of malacophylly = succulence) increases (Skarpe (1996, Lamont et al. 2015). This means that leaf mass area (LMA), the inverse of SLA, may fail as an index of sclerophylly (Groom and Lamont 1999) when malacophylls are included in the study as both are characterized by a marked increase in *z* even though *D*_v_ may stay low. But LMA should still serve as an index of drought tolerance among both leaf types where it implies greater *z* (Lamont & Lamont, 2000; Lamont et al., 2002).

Our collation of data for 115 species that included 33 species designated as succulent by the authors revealed three groups (Fig. 3). The succulent (malacophyll) group was readily identified by its high Q/*A* (= *Qv•z*) associated with a *Q*_m_ of 67−97% and SLA under 20 mm^2^/mg. The overall power-function slope of 0.77 attested to the high *Q*_v_ (minimal air and dry-matter content) and *z* of these species. The mesophyll group by contrast had a *Q*_m_ of 20−88% and SLA up to 45 mm^2^/mg. The overal slope of Q/*A* at 0.59 tended to asymptote at 80% that attested to the moderately high air content and thinness of these species leaves. The high *D*_v_ subgroup with SLA < 10 mm^2^/mg were designated as sclerophylls but did include two of the 34 species that were originally considered as succulent. The latter observation lends support to using the more rigorous criteria of Q/*A* if identification of succulents is one of the objectives of the study. There was also a substantial intermediate group, with a Q/*A* of 0.5−1.0 mg/mm^2^ with limited SLA, that we recognized as semi-malacophylls. The overall pattern from Fig. 3 compared with the equivalent figure in Wang et al. (2022) is that the inclusion of succulents makes the quest for a universal curve quite futile and does not do justice to the rich structural variation in leaf structure that actually exists in nature.

**Fig. 3.**
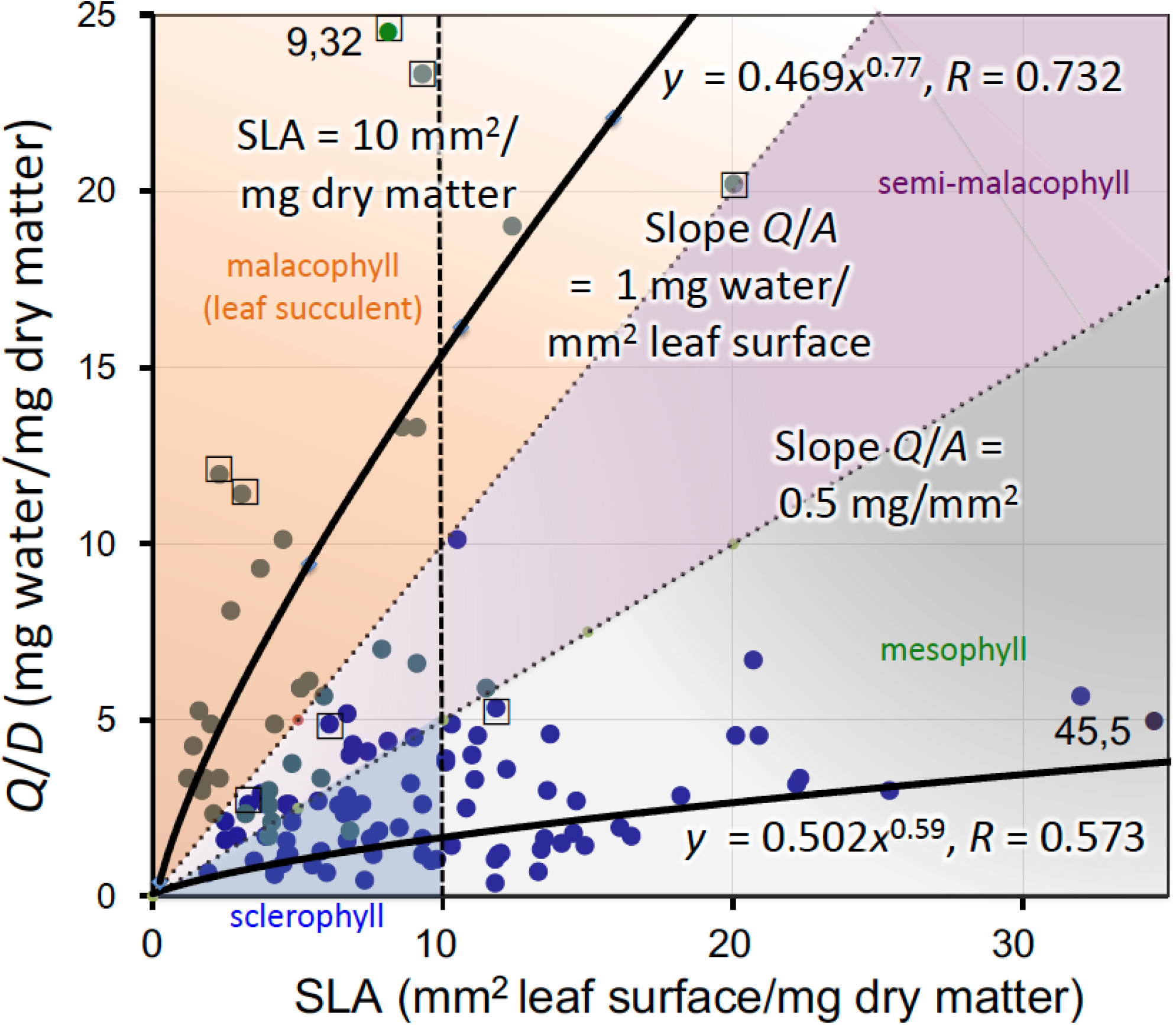
Relationship between *Q*/*D* and SLA among 115 species from this study, Argentina and Spain whose slopes yield *Q*/*A*. Two major groups of species were recognized: high *Q*/*A*, representing succulents (malacophylls, orange triangle)), and low *Q*/*A*, representing sclerophylls (low SLA, blue), and mesophylls (high SLA, grey). An intermediate group of semi-malophylls (pink) was also recognized. Data originally designated as succulents are given as green points, and data give and non-succulents are given as dark blue points. Data points from our study have a square around them. Curve fits were significant at *P* < 0.001.

[We are surprised that so many species had leaves with a *Q*_m_ of 50% or less in Vendramini et al. (2002) and Wang et al. (2022). For our species, bound water ranged 8−27% of mass at saturated water content and levels < 20% were only obtained for the most succulent species (Lamont and Lamont 2000). It seems that saturated water content was often underestimated in these studies, limiting their usefulness for comparative purposes]. More details on comparisons with other studies are given in the Supplementary Information (Appendix).

### Dissolved cations (Na and K)

Our values correspond to species inhabiting the least saline parts of the Negev Desert where (K + Na) is < 5 mmol/kg, assuming 10% dry mass (based on our data) as the results are given on a wet mass basis (Winter *et al*., 1976). *Q*_v_ has a strongly positive but decreasing power relationship with (Na + K) concentration on a dry mass basis (Fig. 4a). But because of the erratic relationhip of *D*_v_ with *Q*_v_ (Fig. S2), the concentration of (K + Na) on a leaf volume basis shows only a tenuous relationship with *Q*_v_ (Fig. 4b). Hence, (K + Na) does not appear to have a critical osmotic function in inducing ‘swelling’ of the vacuoles, as the variation in *Q*_v_ at a given (K + Na) is much too great. Thus, the physiological mechanism responsible for rising succulence must lie elsewhere.

**Fig. 4.**
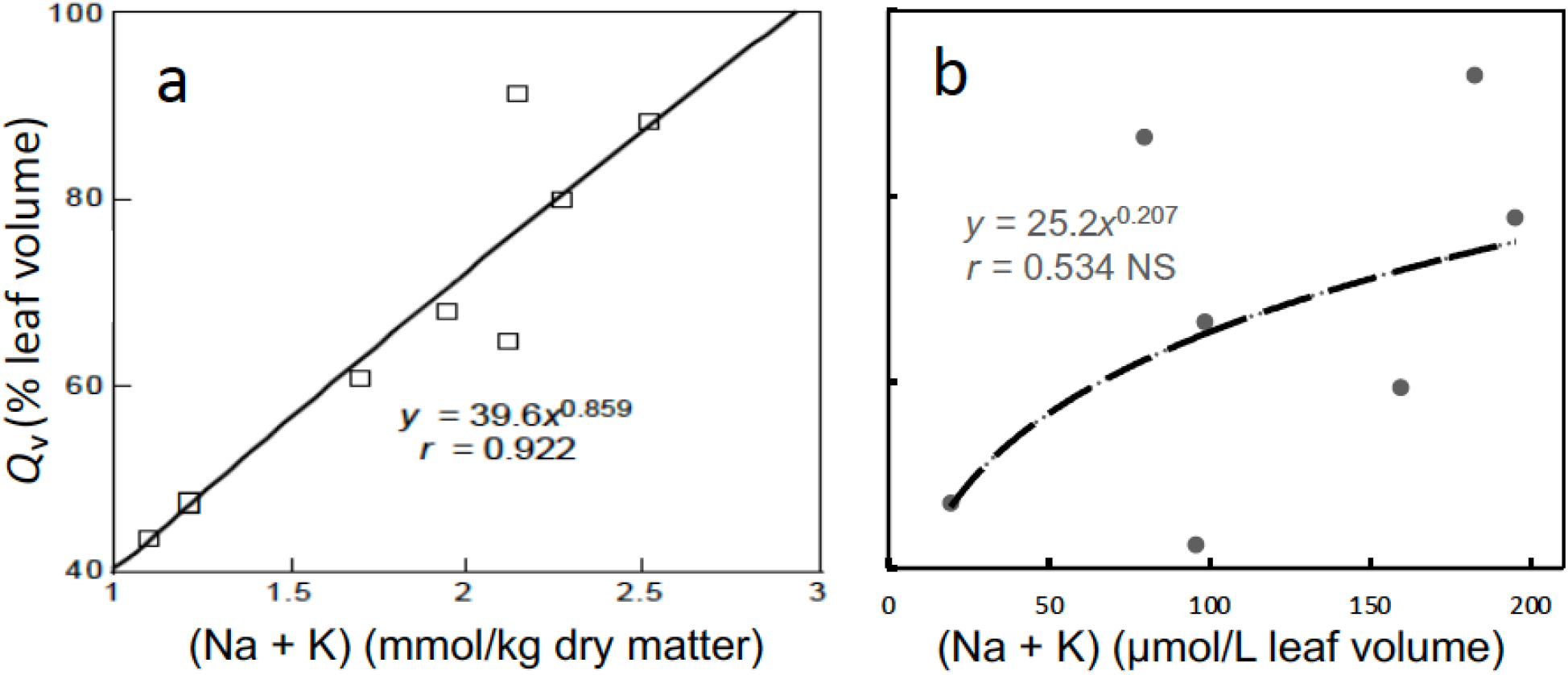
Balancing ions (Na + K) plotted against *Q*_v_, a) on a dry matter baisis, and b) on a leaf volume basis.

### Organic compounds (N and P)

As (*Q* + *D*)/*D*, but not (*Q* + *D*)/*A* and marginally *Q*/*A*, increased, nitrogen on a volume basis (N_v_) decreased in an exponential fashion (Fig. 5a). As *Q*_m_ [= 1 − *D*/(*Q* + *D*), i.e., as (*Q* + *D*)/*D* rises, *Q*_m_ rises] increased, N_v_ decreased strongly in a linear manner while *Q*_v_ was uncorrelated (Fig. 5b). This may be because N is associated with the cytoplasm and hence is included under *D*_m_ (= 1 − *Q*_m_) that decreases as water contributes more to leaf weight. N_m_ was uncorrelated with *Q*_m_ and was omitted from the figure that is difficult to explain as the same argument should have applied. N_v_ and P_v_ were weakly correlated negatively and exponentially with *z* (Fig. 5c) that indicates a strong dilution effect as the leaves thicken and swell with water at the same time. This downward trend agrees with Garnier *et al*. (1999) for N_v_ in grasses but Roderick *et al*. (1999c) obtained no relationship for woody plants. D_m_ was strongly correlated positively with N_v_ and P_v_ (as was *D*_v_ but not shown) as expected from the above rationale (Fig. 5d).

**Fig. 5.**
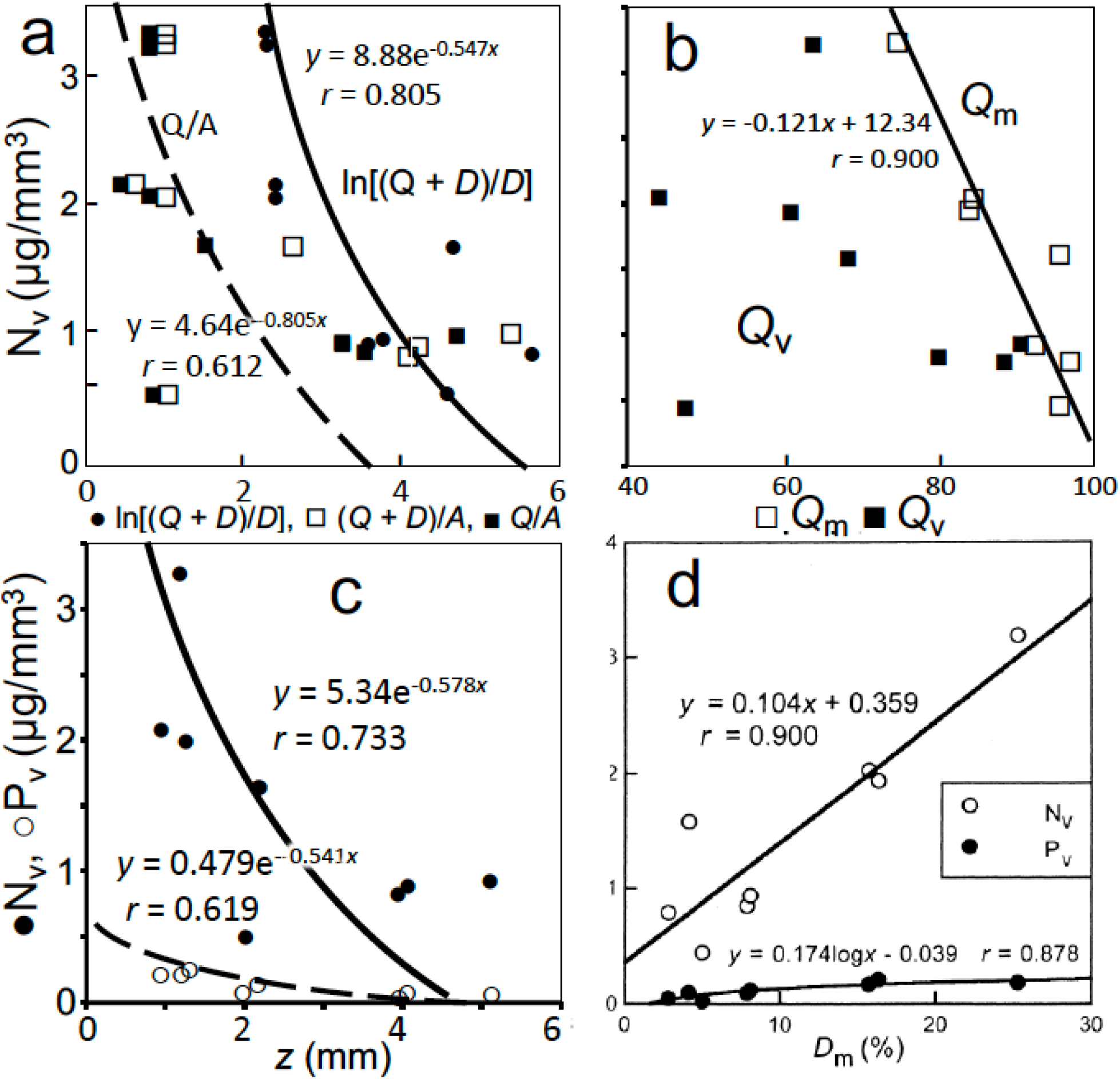
N and P relations. N_v_ plotted against a) (*Q* + *D*)/*D*, (*Q* + *D*)/*A* and *Q*/*A*, and b) *Q*_v_ and *Q*_m_; N_v_ and P_v_ plotted against c) *z* and d) D_m_, all with best-fit lines added.

Generally, there was a dilution effect on N and P as water content on a volume and mass basis increased, or dry matter decreased (Fig. 5). Adding the findings of Roderick *et al*. (1999c), this means that there is no generally applicable relationship between N_m_ and *Q*_m_. Since N resides in the dry matter content of cells, a relationship with *D*_m_ and *D*_v_ seems more likely, at least among succulents (Fig. 5d and unpublished). We also got no relationship between SLA and N_m_ or P_m_ (unpublished), unlike Roderick *et al*. (1999a) but agreeing with Lamont *et al*. (2002) for LMA. In our case, it is probably because *z* and *D*_v_ have opposite effects on N_v_ and P_v_ (Fig. 5c,d). Integrating the results in Figs 3 and 4, cells of a leaf with low succulence (*Qv•z*, e.g., 40% = *Salvia*) possesse relatively smaller vacuoles, larger air spaces (*F*_a_), and slightly thicker walls and higher organic compound content (≡ cytoplasm and nucleus) than one with high succulence (e.g., 90% = *Ruschia*) (Fig. 2).

N on an area basis, N/A, was weakly correlated exponentially with *z*, and P_A_ was best linearly correlated wih *z* (Fig. S3). Since N/A = N_v_•*z*, the weak trend must be due to a tendency for an increase in *z* to override the effects of the associated decrease in N_v_ but more strongly in the case of P/A (Figs. 5c, S3). Garnier *et al*. (1999) also obtained a weak positive relationship between N/A and *z*. This means that the net metabolic activity (photosynthesis, respiration) of the more succulent species would be somewhat more efficient on an area basis, essentially due to their greater *z*, despite the fact that their concentration falls on a volume basis. Lamont et al. (2002), supported by Wright and Westoby (2002), observed a clear drop in N/A and P/A with declining rainfall in their studies that they also correlated with increasing *z*. Thus, increasing drought tolerance, as implied by increasing *z* (Lamont and Lamont 2002), is not accompanied by a loss of metabolic efficiency, despite the dilution of N and P.

### Water-use efficiency (W_i_)

Intrinsic water-use efficiency (*W*_i_, derived from δ^13^C) increased with rising succulence (Q/*A*), increasing by 55% as Q/*A* increased by five times (Fig. 6a). This was essentially due to increasing *z* (Fig. 6b). *W*_i_ halved as SLA increased by ten times (Fig. 6c). There was a non-significant tendency for *D*_v_ and *D*_m_ to decline in a similar way (Fig. 6d) so that SLA’s relationship with *W*_i_ was mainly due to declining *z. W*_i_ showed a tendency to rise linearly with increasing N_A_ but increased significantly with increase in P_A_ (Fig. 6e,f). Since (N,P)/*A* = (N,P)_v_•z, the weak relationship was due to (N,P)_v_ tending to decline as z increased (Fig. 5c) but again the increase must have been due to a dominant *z* effect.

**Fig. 6.**
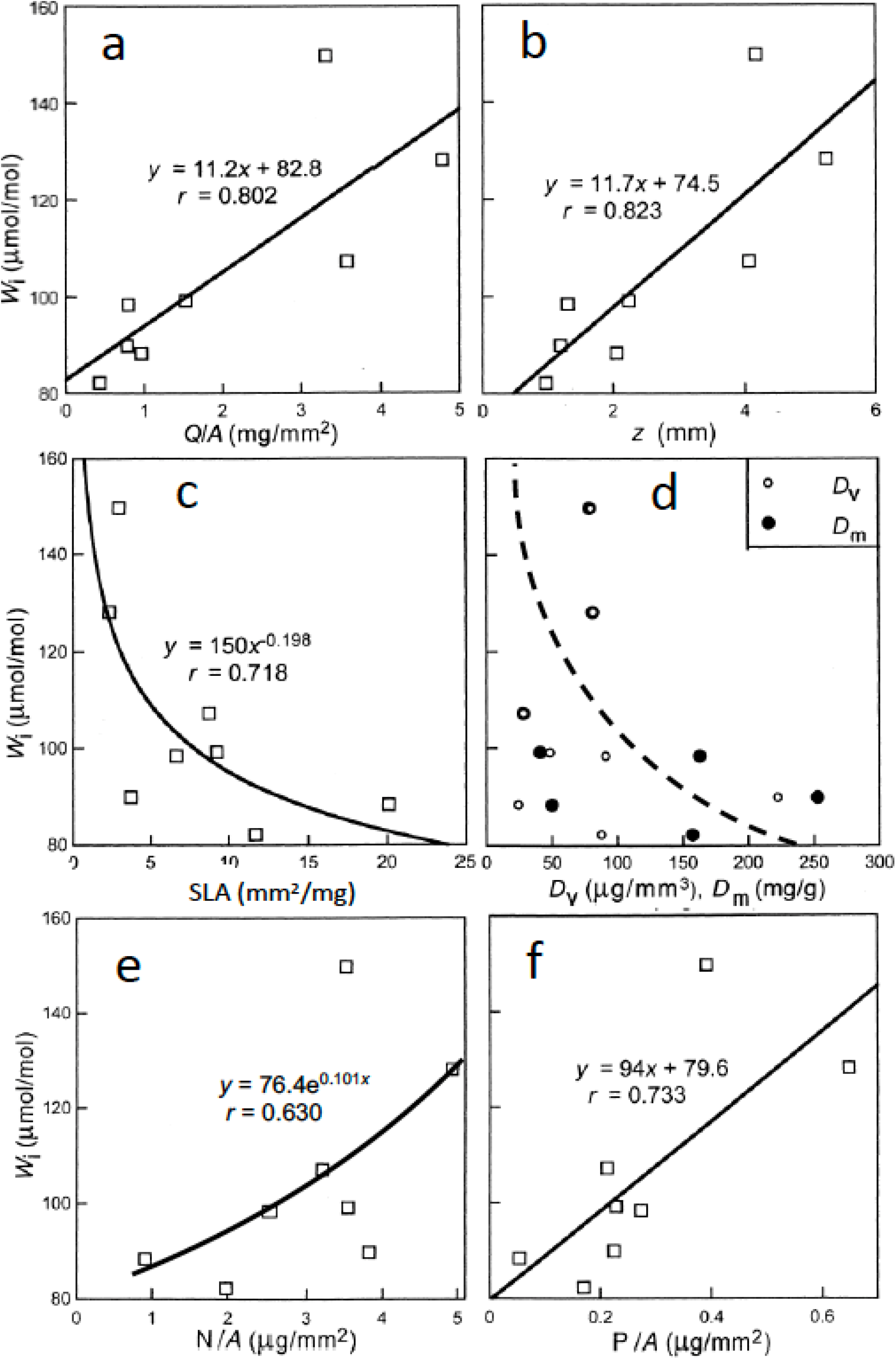
Water-use efficiency (*W*_i_) plotted against a) *Q*/*A*, b) *z*, c) SLA, d) *D*_v_ and *D*_m_, e) N/*A* and f) P/*A*. Best-fit lines added with non-significant trend line dashed.

Our results emphasise that *W*_i_ is a response to both the physiological and structural properties of leaves (Fig. 6). While *Stoeberia* and *Ruschia* (two of the most succulent taxa) have a CAM photosynthetic pathway, the others are C_3_ (Lamont and Lamont 2000), *W*_i_ followed a continuum within and between them. The decline in *W*_i_ with increase in SLA, and rise with increase in *z*, agrees with Schulze *et al*. (1998) and Lamont *et al*. (2002) for sclerophylls. With increasing *z*, stomatal conductance of our species decreases (Lamont & Lamont, 2000), probably controlled by reduced internal conductance as water replaces air space and distances between the xylem and stomata increase (Fig. 2, Hanba *et al*., 1999).

Roderick *et al*. (1999b) noted that, among non-succulents, gas exchange would rise with increase in *z*, no doubt due to the expected increase in N_A_ (Wright & Westoby, 2002). We obtained no clear relationship between *W*_i_ and N_A_ but there was with P_A_. If P limits photosynthesis in this environment then increased photosynthesis may contribute to rising *W*_i_ with increased succulence as well as declining water loss. Both can be attributed to increasing *z* (Figs. 6b, S3). The lack of any relationship between *W*_i_ and dry density (*D*_m_, *D*_v_) indicates that there is no greater demand for carbon per unit area of leaf as succulence increases.

## Conclusions

For a given projected leaf area, *A*, water-storing capacity is dependent on the fraction of volume occupied by water (*Q*_v_) and leaf thickness (*z*). This means that Q/*A* (= *Q*_v_ •*z*) is an ideal index of succulence. SLA cannot be used as an index of succulence as it has no necessary relationship with leaf water or air content. The relationship between SLA and its components and other structural and chemical attributes noted here has parallels with at least some other studies of non-succulent species. The strong positive relationship between *Q*_v_ and *z* has not been noted previously in other groups of plants. Measures of specific gravity (*ρ*_Q+D_ and *ρ*_l_) have similar values to those for mesophytes-sclerophylls but they have a different relationship with the air space (*F*_a_), highlighting differences in control of *ρ*_l_ between the two groups. Rising succulence is accompanied by a dilution in the concentration of N and P but a rise in Na and K. It appears that *z* has a key role in controlling *W*_i_. When our data are combined with (the few) other studies that included leaf succulents it is clear from *Q*/*A* that there is a completely different relationship between *Q*_m_ and SLA among succulents (malacophylls) and sclerophyll-mesophylls so that a universal curve is neither possible nor desirable, despite recent attempts (Wang et al. 2022). High leaf thickness, *z*, holds the key to the special physiological and structural features of leaf succulents. Since succulence is a structural concept, *Q*_v_ (water content on a volume basis) is a much more useful parameter than *Q*_m_ (water content on a mass basis) in understanding its relationship with other variables.

While some clear trends emerged in our study, the sample size was small (although the range of values obtained attests to our great success in selecting species to cover previous extremes, often greatly exceeding them, Table 1) and our measurements need to be repeated on other species in other dry regions that vary in degree of succulence.

## Abbreviations

δ^13^C: discrimination ratio between ^12^C and ^13^C
*ρ*_l_: leaf specific gravity (*D*_v_ compared with mass of an equal volume of water)
*ρ*_Q+D_: specific gravity on a mass basis (mass of water at saturation and dry matter compared with mass of water with the same volume as the water and dry matter)
*Ψ*: water potential
*A*: leaf projected area
*D*: dry mass per leaf
*D*_m_: dry matter on a leaf mass basis
*D*_v_: dry matter on a leaf volume basis
*F*_a_: fraction of leaf volume as air space
*F*_D_: fraction of leaf volume as dry matter
l: referring to the leaf
m: referring to leaf mass
m: referring to mass
LMA: the inverse of SLA
N_m_: nitrogen content on a dry mass basis
N_v_: nitrogen content on a volume basis
P_m_: phosphorus content on a dry mass basis
P_v_: phosphorus content on a volume basis
SLA: specific leaf area (*A*/*D* = 1/*D*_m_•z)
*Q*: water mass per leaf
*Q*_m_: water content on a dry mass basis
*Q*_v_: water content on a volume basis (= *F*_Q_)
v: referring to leaf volume
*V*: volume per leaf
*V*_D_: dry matter volume per leaf
*W*_i_: intrinsic water-use efficiency (based on δ^13^C)
*z*: adjusted leaf thickness (after Lamont et al. 2015).

## Supplementary information

Fig. S1. Three standard indices of leaf succulence plotted against SLA. Fig. S2. *Q*_v_ plotted against *D*_v_.

Fig. S3. Relationship between N/*A*, P/*A* and *z*. Fig. S4. Relationship between *D*_v_ and *D*_m_.

Appendix: Further comparisons of SLA with those in the literature.

## Acknowledgements

This work was undertaken in the Department of Botany, University of Stellenbosch, Western Cape, South Africa with the assistance of a FRD Research Fellowship to Byron Lamont that was otherwise self-funded. We thank Karen Esler, Neil Eccles, Mike Cramer, Valdon Smith and Alex Valentine for their invaluable help, and also Ian Wright, Michael Roderick and Philip Groom for their insightful comments on a draft of the manuscript. Data will be made available on acceptance of the manuscript.

## Author contributions

BBL conceived, managed, undertook most field and laboratory work, liaised with other researchers, performed statistics, prepared the figures, wrote the manuscript and oversaw the publication process. HCL participated in all field and laboratory work and approved of the submitted manuscript.

